# On the feasibility of saltational evolution

**DOI:** 10.1101/399022

**Authors:** Mikhail I. Katsnelson, Yuri I. Wolf, Eugene V. Koonin

## Abstract

One of the key tenets of Darwin’s theory that was inherited by the Modern Synthesis of evolutionary biology is gradualism, that is, the notion that evolution proceeds gradually, via accumulation of “infinitesimally small” heritable changes ^1,2^. However, some of the most consequential evolutionary changes, such as, for example, the emergence of major taxa, seem to occur abruptly rather than gradually, as captured in the concepts of punctuated equilibrium ^3,4^ and evolutionary transitions ^5,6^. We examine a mathematical model of an evolutionary process on a rugged fitness landscape ^7,8^ and obtain analytic solutions for the probability of multi-mutational leaps, that is, several mutations occurring simultaneously, within a single generation in one genome, and being fixed all together in the evolving population. The results indicate that, for typical, empirically observed combinations of the parameters of the evolutionary process, namely, effective population size, mutation rate, and distribution of selection coefficients of mutations, the probability of a multi-mutational leap is low, and accordingly, their contribution to the evolutionary process is minor at best. However, such leaps could become an important factor of evolution in situations of population bottlenecks and elevated mutation rates, such as stress-induced mutagenesis in microbes or tumor progression, as well as major evolutionary transitions and evolution of primordial replicators.

---

Within the framework of modern evolutionary biology, gradualism corresponds to the weak-mutation limit, that is, an evolutionary regime in which mutations occur one by one, consecutively, such that the first mutation is assessed by selection and either fixed or purged from the population, before the second mutation occurs ^9^. An opposite, saltational mode of evolution ^10,11^ is imaginable under the strong-mutation limit ^9^ whereby multiple mutation occurring within a single generation and in the same genome potentially could be rejected or fixed all together. Under the fitness landscape concept ^7,8^, gradual or more abrupt evolutionary processes can be depicted as distinct types of moves on rugged fitness landscapes (Figure 1). The typical evolutionary paths on such landscapes are thought to be one step at a time, uphill mutational walks ^8^. In small populations, where genetic drift becomes an important evolutionary factor, the likelihood of downhill movements becomes non-negligible ^12^. One can imagine, however, a radically different type of moves on these landscapes, namely, leaps (or “flights”) across valleys when a population can move to a different area in the landscape, for example, to the slope of a different, higher peak, via simultaneous fixation of multiple mutations (Figure 1).

**Figure 1.**
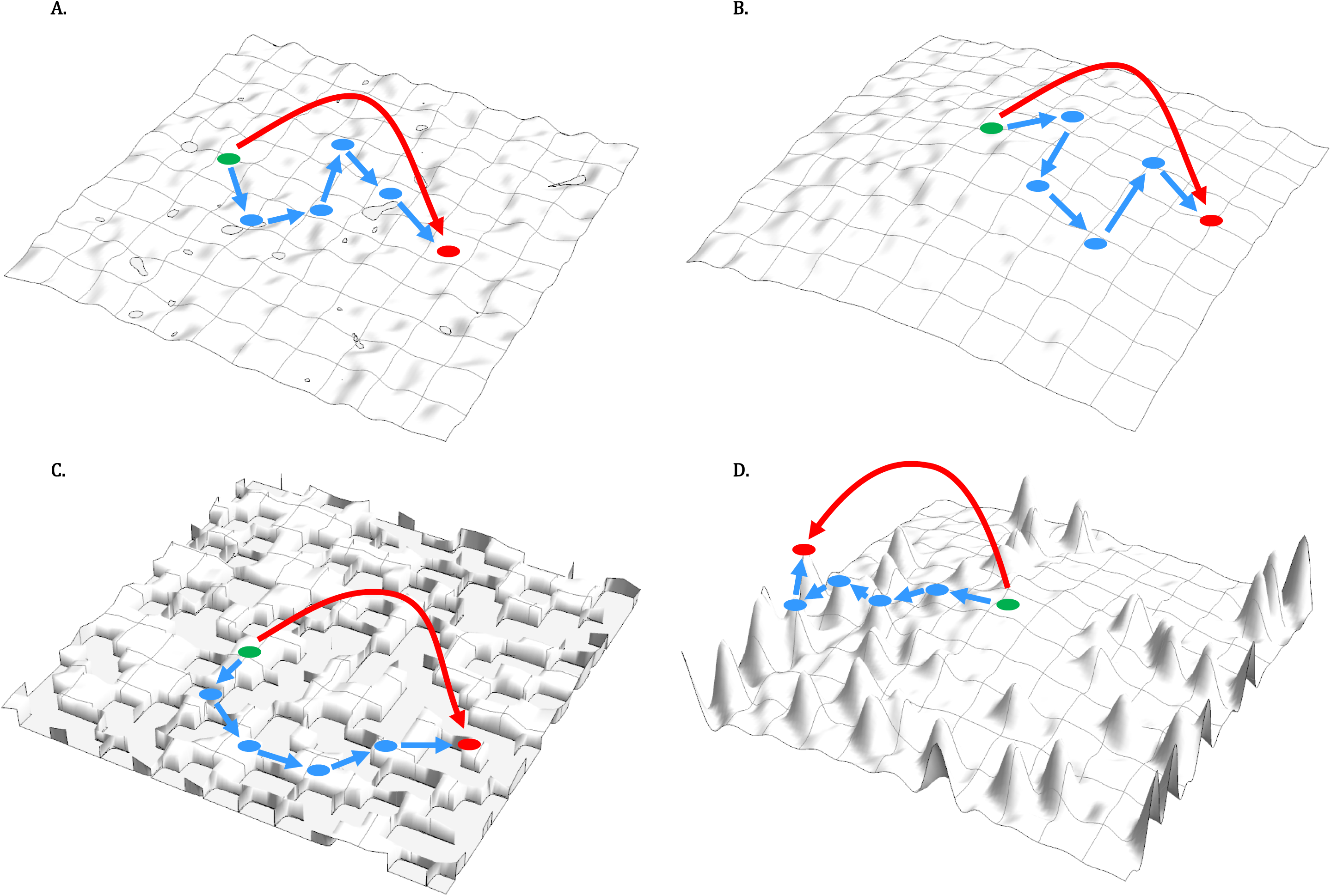
Walks and leaps on different types of fitness landscapes. Dots show genome states; blue arrows indicate consecutive moves via fixation of single mutations; red arrows indicate multi-mutational leaps. A. Nearly neutral landscape. B. Landscape dominated by slightly deleterious mutations. C. Kimura’s model landscape (a fraction of mutations is neutral; the rest are lethal). D. Landscape combining beneficial and deleterious mutations.

We sought to obtain analytically, within the population genetics framework, the conditions under which such leaps might be feasible. Let us assume (binary) genomes of length *L* (in the context of this analysis, *L* should be construed as the number of evolutionarily relevant sites, such as codons in protein-coding genes, rather than the total number of sites), the probability of single mutation *μ* << 1 per site per round of replication (generation), and effective population size *N* >> 1. Then, the transition probability from sequence *i* to sequence *j* is (Ref. 13, Eq.3.11):

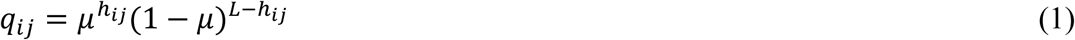

where *h*_*ij*_ is the Hamming distance (number of different sites between the two sequences). The number of sequences separated by the distance *h* is equal to the number of ways *h* sites can be selected from *L*, that is,

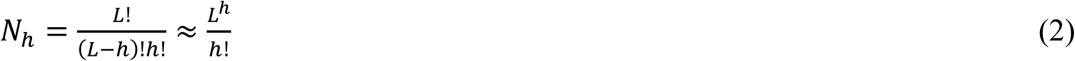

where the last, approximate expression is valid assuming that *L* >> 1 and *L* >> *h* (*h* can be of the order of 1).

Assuming also *μ* << 1, we obtain a typical combinatorial probability of leaps at the distance *h*:

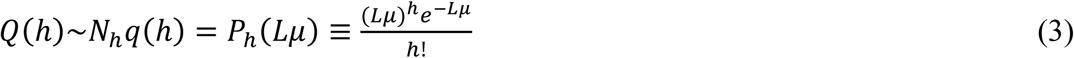

which is a Poisson distribution with the expectation *Lμ*.

At the steady state, the probability of fixation of the state *i* is proportional to exp(-*vx*_*i*_) where

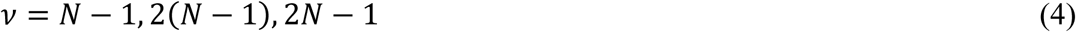

for the Moran process, haploid Wright-Fisher process, and diploid Wright-Fisher process, respectively, and *x*_*i*_ = - ln *f*_*i*_ where *f*_*i*_ is the fitness of the genotype *i* (*x*_*i*_ is analogous to energy in the Boltzmann distribution within the analogy between population genetic and statistical physics ^14^). Then, the rate of the occurrence and fixation of the transition *i* → *j* is ^13^

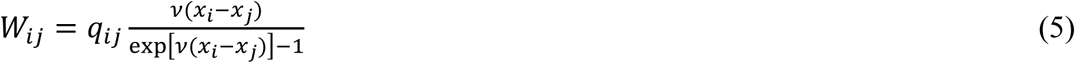

The distribution function of the fitness difference Δ_*ij*_ = *x*_*i*_ - *x*_*j*_ has to be specified (hereafter, we refer to *x* as fitness, omitting logarithm for brevity). We analyze the case without epistasis, that is, with additive fitness effects of individual mutations:

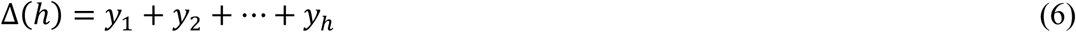

where *y*_*i*_ are independent random variables with the distribution functions *G*_*j*_(*y*_*j*_). Then, the distribution function of the fitness difference is

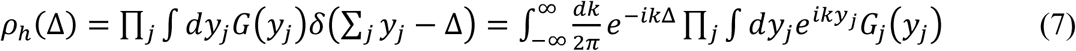

which is obtained by using the standard Fourier transformation of the delta-function.

Now, let us specify the distribution of the fitness effects of mutations *G*_*j*_(*y*_*j*_), assuming an exponential dependency of the probability of fixation of a mutation on its fitness effect, separately for beneficial and deleterious mutations:

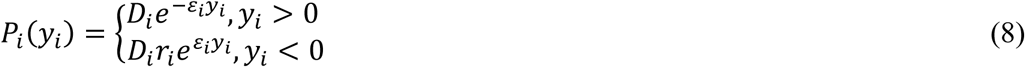

where *D*_*i*_ is the normalization factor, *r*_*i*_ is the ratio of the probabilities of beneficial and deleterious mutations, and *ε*_*i*_ is the inverse of the characteristic fitness difference for a single mutation (see below). For simplicity, we assume here the same decay rates for the probability density of the fitness effects of beneficial and deleterious mutations. Empirical data on the distributions of fitness effects of mutations ^15,16^ clearly indicate that *r*_*i*_ ≪ 1. From the normalization condition,

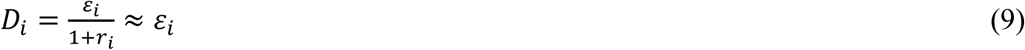

Note that the mean of the fitness difference (selection coefficient) when the distribution of the fitness effects is given by (8) is

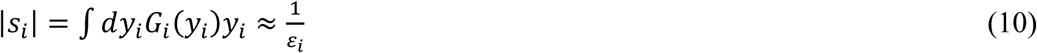

For simplicity, we start with an assumption that the values of *D*_*i*_ and *r*_*i*_ are independent of *i*. For the model (8):

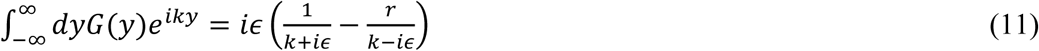

Then, from equation (5), the rate of fixation is equal to

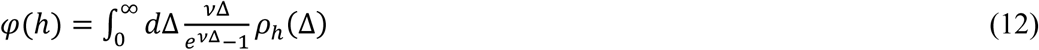

Substituting (11) to (7), we obtain

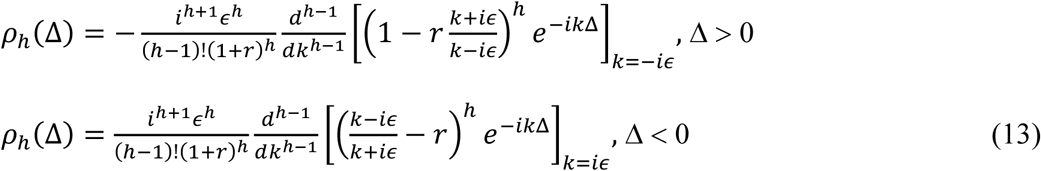

Consider first the case *r* = 0 (all mutations are deleterious). Then, *ρ*_*h*_(Δ < 0). For Δ > 0, that is, decrease of the fitness, we have:

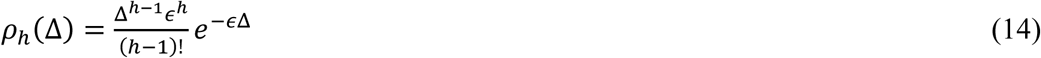

Then, the fixation rate (12) of a leap at a distance *h* is equal to

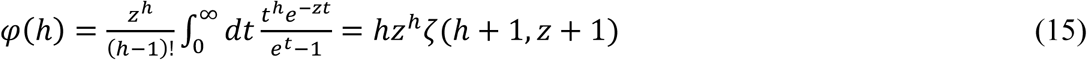

where *z* = *є*/*v* and *ζ*(*x,y*) is the Hurwitz zeta function 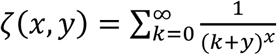. Therefore, the rate of fixed leaps of the length *h* is equal to *W*(*h*) = *P*_*h*_(*Lμ*) *φ*(*h*).

In one extreme, if *z* ≫ 1 (*v* | *s* |≪ 1, neutral landscape), *φ*(*h*) ≈ 1 and mutations are fixed at the rate they occur. In the opposite extreme case of strong negative selection (*z* ≪ 1, *v* | *s* |≫ 1), *φ*(*h*) ≈ *hz*^*h*^ *ζ*(*h* + 1) where *ζ*(*x*) is the Riemann zeta function. For a rough estimate, *ζ*(*h* + 1) can be replaced by 1, and then, *W(h*) ≈ *Lμe*^−*Lμ*^*P*_*h* − 1_ (*Lμz*). In this case, the maximum of *W(h*) is reached at 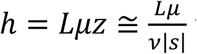 which gives a non-negligible fraction of multi-mutation leaps (*h* > 1) among the fixed mutations only for *Lμ* ≥ *v* | *s* |. However, in this case, the value of *W(h*) at this maximum is exponentially small because *e*^−*Lμ*^ < *e*^−*v*| *s* |^. Therefore, in the regime of strong selection against deleterious mutations and at high mutations rates (*Lμ* ≥ *v* | *s* |), multiple mutations actually dominate the mutational landscape, but their fixation rate is extremely low. Qualitatively, this conclusion seems obvious, but we now obtain the quantitative criteria for what constitutes “strong selection”. We find that, even for *v*|*s*| ∼ 10, the rate of multi-mutation leaps (*h* = 4) can be non-negligible (>10^-4^ per generation, Figure 2A) at the optimal *Lμ* values, whereas for *v*|*s*| ∼ 100, any leaps with *h*>1 are unfeasible (Figure 2B).

**Figure 2.**
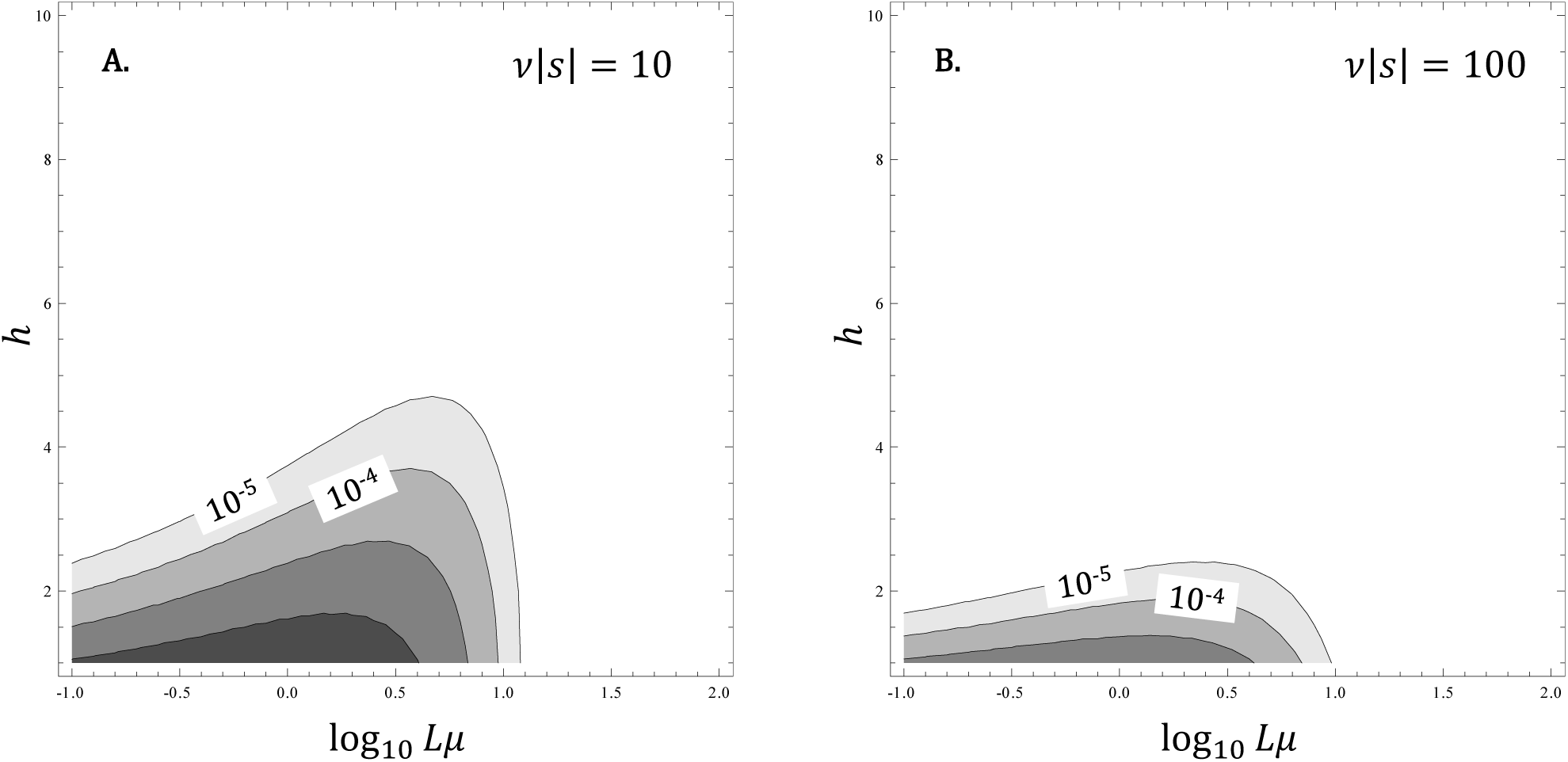
Rates of leaps on a landscape dominated by deleterious mutations. Rates of transitions are plotted against the per-genome mutation rate (*Lμ*) and the leap length for different strengths of selection (A: ν|*s*| = 10 and B: ν|*s*| = 100). Contour lines indicates orders of magnitude and start from the rate of 10^-5^ leaps per generation.

Under a more realistic model, all values of *ε*_*i*_ (fitness effects of mutations) are different. For Δ > 0 and *r* = 0 (no beneficial mutations)

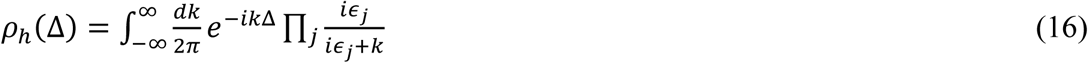

For example, in Kimura’s neutral evolution model^17^, *ε*_*i*_ is a binary random variable that takes a value of ∞ (|*s*_*i*_| = 0, neutral mutation), with the probability *f*, and a value of 0 (|*s*_*i*_| = ∞, lethal mutation), with the probability 1 - *f*. Then, *ρ*_*h*_ (Δ), = *f*^*h*^*δ* (Δ), *φ*(*h*) = *f*^*h*^ and *Lμ* is replaced with *Lfμ* in equation (3), a trivial replacement of the total genome length *L* with the length of the part of the genome where mutations are allowed, *Lf*. Accordingly, *W*(*h*) = *P*_*h*_(*Lfμ*), and multi-mutational leaps become relevant for *Lfμ* ≥ 1.

Let us now estimate the probability of leaps with beneficial mutations (Δ < 0). Assuming *r* ≪ 1(rare beneficial mutations), equation (13) takes the form

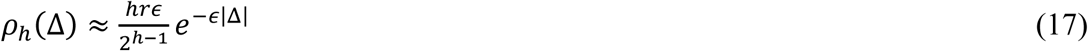

and the fixation rate of beneficial mutations is

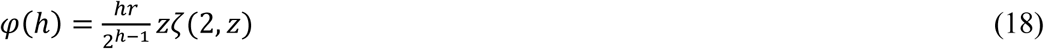

If *z* ≫ 1 (weak positive selection), *z ζ*(2, *z*) ≈ 1, so that the role of beneficial mutations is negligible. If *z* ≪ 1 (strong positive selection),

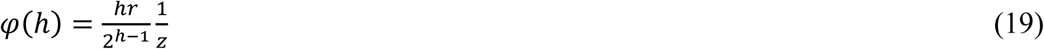

Comparing equation (19) with the result for Δ > 0 (equation (18)), one can see that, in this case, beneficial mutations are predominant among the fixed mutations if

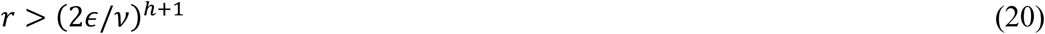

In this regime, multi-mutation leaps (*h* > 4), occur at non-negligible rates under sufficiently high (but not excessive) mutation rates (Figure 3).

**Figure 3.**
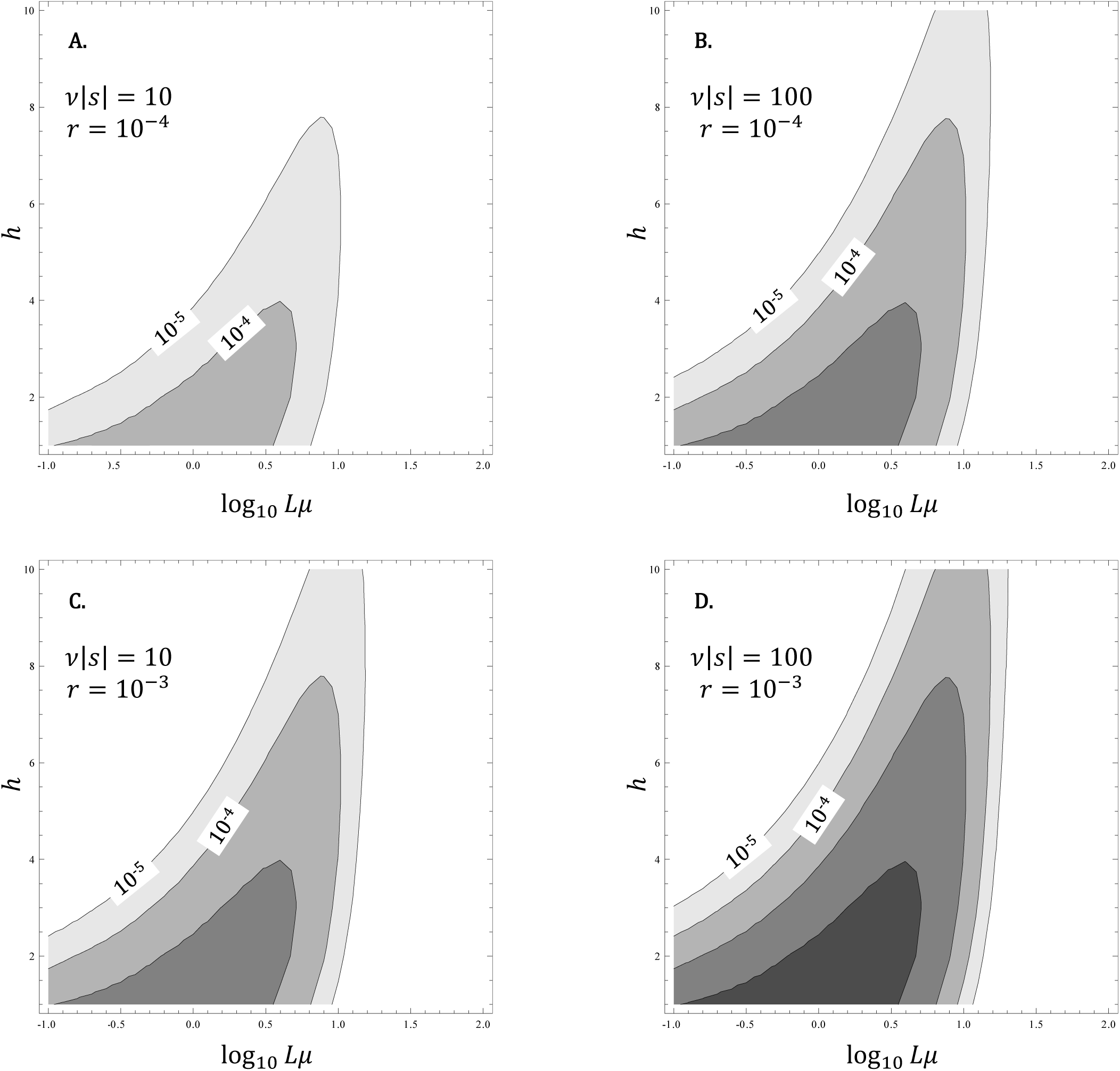
Rates of leaps on a landscape combining beneficial and deleterious mutations. Rates of leaps are plotted against the per-genome mutation rate (*Lμ*) and the leap length for different strengths of selection (A and C: ν|*s*| = 10; B and D: ν|*s*| = 100) and for different frequencies of beneficial mutations (A and B: *r* = 10^-4^; C and D: *r* = 10^-3^). Contour lines indicates orders of magnitude and start from the rate of 10^-5^ leaps per generation.

Note that, throughout this analysis, we only address simple cases of evolution without epistasis. Including the more realistic situation with epistasis will be the direction of further development of these models. Likewise, we disregard possible effects of clonal interference when calculating the mutation fixation probability.

In summary, we obtained analytic expressions for the probability of multi-mutational leaps for deleterious and beneficial mutations depending on the parameters of the evolutionary process, namely, effective genome size (*L*), mutation rate (*μ*), effective population size (*v*), and distribution of selection coefficients of mutations (*s*). Leaps in random fitness landscapes in the context of punctuated equilibrium have been previously considered for infinite^18,19^ or finite^20^ populations. However, unlike the present work, these studies have focused on the analysis of the dynamics of the leaps rather than on the equilibrium distribution of their lengths.

The principal outcome of the present analysis are the conditions under which multi-mutational leaps occur at a non-negligible rate in different evolutionary regimes. If the landscape is completely flat (strict neutrality, *s* = 0), the leap length is distributed around *Lμ*, that is simply, the expected number of mutations per genome per generation. If *Lμ* ≪ 1, leaps are effectively impossible, and evolution can proceed only step by step, under the “Russian roulette” model on a graph with edges linking only neighboring nodes ^8^. A considerable body of data exists on the values of each of the relevant parameters that define the probability of leaps. Generally, in the long term, the total expected number of mutations per genome per generation has to be of the order of 1 or lower because, if *Lμ* ≫ 1, the population ultimately succumbs to mutational meltdown ^13,21^. The selection for lower mutation rates is thought to be limited by the drift barrier and, accordingly, the genomic mutation rate appears to be inversely proportional to the effective population size, that is, *Lμ* ∼ 1/ *v*^22,23^. Thus, *Lμv* = *const,* apparently an important universal in evolution. Under this “law”, organisms with small effective population sizes, for example, large mammals cannot evolve low mutation rates and exist closer to the mutational meltdown threshold than organisms with large populations, such as bacteria ^23^.

To estimate the leap probability, we can use equation (15) and characteristic values of the parameters, for example, those for human populations. As a crude approximation, *Lμ* = 1*,v* = 10^4^, |*s*| = 10^-2^ which, in the absence of beneficial mutations, translates into the probability of a multi-mutation leap of about 4×10^-5^. Thus, such a leap would, on average, require over 23,000 generations which is not a relevant value for the evolution of mammals (given that ∼140 single mutations are expected to be fixed during that time as calculated using the same formula).

However, short leaps comprised of beneficial mutations can occur with reasonable rates, such as 5×10^-4^for *h* = 3, and the frequency of beneficial mutations of *r* = 10^-4^, and such leaps are only 8 times less frequent than single mutation fixations. Conceivably, such leaps of beneficial mutations could be a minor but non-negligible evolutionary factor. For organisms with *Lμ* < 1 and larger *v,* the probability of leaps is lower than the above estimates, so that under “normal” evolutionary regimes, the contribution of leaps is small at best.

However, in some biologically relevant and common situations, such as stress-induced mutagenesis, which occurs in microbes in response to double-stranded DNA breaks, the effective mutation rate can locally and temporarily increase by orders of magnitude ^24,25^ while the population is going through a severe bottleneck. Roughly, for such a case, we can assume *Lμ* > 1*, v* = 10^3^, |*s*| = 10^-2^. Then, the highest rate of leaps, even in the absence of beneficial mutations, is about 5×10^-3^ (once every 200 generations) and is observed at *Lμ* ≈ 2 (Figure 2A). For beneficial mutations, even longer leaps (e.g., *h* = 7) can occur at non-negligible rates, on the order of 1.5×10^-4^(one leap per 5,000-10,000 generations) if both the mutation rate and selection are sufficiently strong (*Lμ* = 8, *r* = 10^-4^, *v*|*s*| = 100; under these parameters, multi-mutational leaps outnumber single mutation fixations by a factor of ∼50). Thus, it appears that leaps can be an important factor of adaptive evolution under stress.

More generally, severe population bottlenecks that can be caused by environmental stress, and especially, catastrophic events, are known as times of evolutionary innovation when slightly or moderately deleterious mutations that are weeded out by purifying selection in larger populations can be fixed via genetic drift ^12,26^-^28^. The findings reported here show that the innovation potential ^29^ of population bottlenecks could be even higher than considered previously thanks to the possibility of abrupt major changes brought about my multi-mutational leaps. Such leaps can be one of the keys to the puzzle of the evolution of complex adaptations requiring multiple mutations that are adaptive all together whereas each is neutral or even slightly deleterious on its own ^11,30^-^32^. Evolution of such adaptations is a long-standing challenge for evolutionary theory that goes back to Darwin’s discussion of the evolution of the eye ^1^ and sometimes summons the specter of “irreducible complexity” ^33,34^. The leaps could partially overcome the obstacles on the evolutionary path to complexity and might be particularly impactful at times of major evolutionary transitions that likely involve severe stress and extreme population bottlenecks ^6,35^. Another area where leaps potentially could be important is tumor evolution. In many tumors, the effective population size, that is, the number of actively proliferating cells, such as cancer stem cells, is small whereas the mutation rate is high ^36^. Again, this is a situation that is particularly conducive to leaps, and it cannot be ruled out that combinations of 2 to 5 driver mutations that are typically required for malignant transformation ^37^-^40^, in some tumors, could be fixed in a single leap.

Finally, primordial replicators, in particular, those in the hypothetical RNA World, are thought to have had an extremely low replication fidelity, barely above the mutational meltdown threshold ^41^-^43^. Under these conditions, leaps could have been an important route of evolutionary acceleration and thus could have contributed substantially to the most challenging evolutionary transition of all, that from pre-cellular to cellular life forms.

Taken together, all these biological considerations suggest that multi-mutation leaps, especially, those including beneficial mutations, the probability of which we show to be non-negligible under conditions of a population bottleneck accompanied by elevated mutagenesis, could be an important mechanism of evolution that so far has been largely overlooked. Saltational evolution, after all, might substantially contribute to the history of life, and in particular, to the emergence of complexity, in direct defiance of the “*Natura non facit saltus*’ principle.

